# Illuminating the Dark Cancer Phosphoproteome Through a Machine-Learned Co-Regulation Map of 26,280 Phosphosites

**DOI:** 10.1101/2024.03.19.585786

**Authors:** Wen Jiang, Eric J. Jaehnig, Yuxing Liao, Tomer M. Yaron-Barir, Jared L. Johnson, Lewis C. Cantley, Bing Zhang

## Abstract

Mass spectrometry-based phosphoproteomics offers a comprehensive view of protein phosphorylation, but limited knowledge about the regulation and function of most phosphosites restricts our ability to extract meaningful biological insights from phosphoproteomics data. To address this, we combine machine learning and phosphoproteomic data from 1,195 tumor specimens spanning 11 cancer types to construct CoPheeMap, a network mapping the co-regulation of 26,280 phosphosites. Integrating network features from CoPheeMap into a machine learning model, CoPheeKSA, we achieve superior performance in predicting kinase-substrate associations. CoPheeKSA reveals 24,015 associations between 9,399 phosphosites and 104 serine/threonine kinases, including many unannotated phosphosites and under-studied kinases. We validate the accuracy of these predictions using experimentally determined kinase-substrate specificities. By applying CoPheeMap and CoPheeKSA to phosphosites with high computationally predicted functional significance and cancer-associated phosphosites, we demonstrate the effectiveness of these tools in systematically illuminating phosphosites of interest, revealing dysregulated signaling processes in human cancer, and identifying under-studied kinases as putative therapeutic targets.

## Introduction

Protein phosphorylation is a crucial post-translational modification (PTM) that regulates a wide range of cellular processes, such as proliferation, differentiation, motility, and apoptosis^1^. This modification is tightly controlled by protein kinases and phosphatases^2^. Dysregulated protein phosphorylation, often driven by aberrant kinase activity, is implicated in many diseases, including cancer^3–5^. Consequently, kinases have become a promising class of drug targets for developing cancer therapies, with more than 70 kinase inhibitors approved for cancer treatment since the approval of imatinib for chronic myeloid leukemia in 2001^6^. However, these inhibitors only target 50 out of the more than 500 protein kinases encoded by the human genome, leaving the majority of kinases open for further exploration and investigation.

Advancements in proteogenomics have opened new horizons in understanding the molecular intricacies of cancer biology^7–20^. Central to these advancements is the exploration of phosphoproteomic landscapes using mass spectrometry (MS)-based phosphoproteomics, which has transformed the analysis of protein phosphorylation, enabling comprehensive, unbiased profiling of phosphorylation events across the entire proteome^21^. Its application in cancer research has been growing rapidly due to the central role of phospho-signaling in cancer biology and treatment^22^. A single phosphoproteomic investigation can uncover tens of thousands of phosphosites, with quantitative comparisons often revealing hundreds to thousands of significantly regulated sites. Annotating phosphosites with existing functional and regulatory information is crucial for interpreting phosphoproteomics data. However, this presents a significant challenge, as less than 5% of the human phosphoproteome has been experimentally linked to specific kinases, and even fewer phosphosites have been functionally characterized. The paucity of knowledge on phosphosites has led to the notion of the “dark phosphoproteome”^23^.

A common task in phosphoproteomic data analysis is to infer upstream kinase activity changes based on phosphoproteomic data. This analysis relies on known kinase-substrate associations (KSAs), excluding the vast majority of phosphosites without annotated kinases. Many computational methods have been developed to predict KSAs^24^, but most utilize consensus sequence motifs or position-specific scoring matrices derived from known KSAs for individual kinases. Because 90% of the annotated phosphosites are associated with just 20% of kinases that are the most well-studied, our ability to predict KSAs and infer activities for the under-studied kinases is restricted, despite their potential as new therapeutic targets.

Biological networks, such as protein-protein interaction networks and gene co-expression networks, are effective means of propagating information from well-characterized molecular components to under-studied ones, leading to a better understanding of their regulation and functions^25^. These networks have proven successful in predicting functions for under-studied proteins^26^ and identifying new disease genes^27^. Because one kinase can regulate multiple phosphorylation events, and co-regulated phosphosites are more likely to participate in the same biological processes, we reason that a co-regulation network of phosphosites, even in the absence of specific knowledge about their regulatory kinases, can provide valuable insights into the functional characterization and regulatory mechanisms of the dark phosphoproteome.

In this paper, we utilize machine learning and phosphoproteomic data from 1,195 clinical tumor specimens spanning 11 cancer types recently harmonized by the National Cancer Institute’s Clinical Proteomic Tumor Analysis Consortium (CPTAC) to construct CoPheeMap, a co-regulation network of 26,280 phosphosites. By incorporating network embedding features from CoPheeMap into a machine learning model called CoPheeKSA, we achieve superior performance in predicting KSAs, uncovering 24,015 novel associations, many of which involve under-studied kinases. Furthermore, we validate these CoPheeKSA predictions by comparing them to substrate specificities determined experimentally using the recently published Kinase Library (KL) approach^28^. To illustrate the practical utility of CoPheeMap and CoPheeKSA, we present examples demonstrating their ability to systematically shed light on phosphosites of interest, revealing dysregulated signaling processes in human cancer, and identifying under-studied kinases as putative therapeutic targets.

## Results

### Phosphosite co-expression is an effective predictor of co-regulation

This study utilized recently harmonized pan-cancer (PanCan) proteogenomic data from the CPTAC pan-cancer resource working group^8^. The harmonized dataset contained phosphoproteomic data for 1,195 tumor samples across 11 cancer types, and phosphoproteomic data was also available for normal samples of 9 cancer types (**Figure 1A**). In total, the dataset covered 158,796 phosphosites, including 77,442 quantified in at least 20% of samples in one tumor cohort. Global proteomic and RNASeq data were also available for the same cohorts, covering a total of 14,103 and 21,592 protein-coding genes, respectively (**Figure 1A**). Among the quantified phosphosites, fewer than 5% were annotated with up-stream kinases or known functions (**Figure 1B**), echoing the widely recognized dark phosphoproteome challenge^23^.

**Figure 1:**
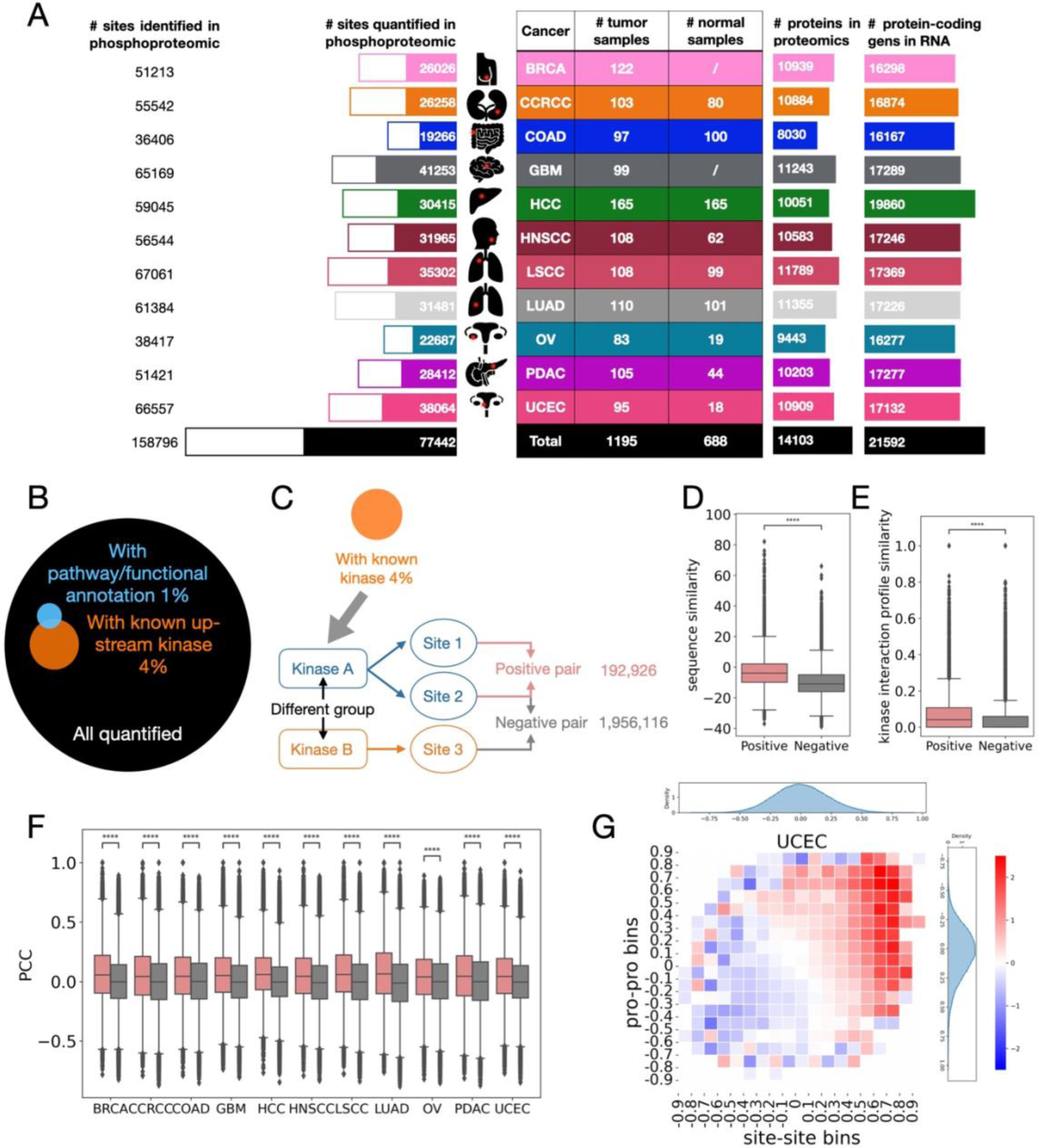
PanCan datasets and ground truth of co-regulated site pairs. A) Diagram indicating numbers of tumor/normal samples, of phosphosites identified in total and quantified in at least 20% samples, and of genes measured by proteomics and transcriptomics in a cohort of 11 cancer types (BRCA, CCRCC, COAD, GBM, HCC, HNSCC, LSCC, LUAD, OV, PDAC and UCEC). A summary of proteomics and transcriptomics from 11 cancer types. B) > 95% of the phosphosites in the PanCan datasets are dark. C) Ground truth data including 192,926 unique positive (targets of the same kinase) and 1,956,116 negative (targets of kinases from different groups) site pairs. D-F) Ground truth co-regulated phosphosites have significantly higher sequence similarity scores, kinase interaction profile similarity scores and site-site correlations compared to the negative pairs. G) 2-dimensional Log Likelihood Ratio (LLR) plot for UCEC. Site-site abundance correlations are shown on the x-axis. Protein-protein abundance correlations are shown on the y-axis.

We reasoned that phosphosites co-regulated by the same kinase are likely to be correlated with each other across different tumors because the set of target sites of that kinase is expected to show higher levels of phosphorylation in tumors with higher kinase activity than those with lower kinase activity. To formally assess the relationship between phosphosite co-expression and co-regulation, we constructed a “ground-truth” dataset of phosphosite co-regulation. This dataset was created from a previously published database comprising comprehensively curated 14,679 KSAs (352 kinases, 9,526 unique sites) from the literature^29^. Among those KSAs, 4,873 KSAs involved the phosphosites quantified in our PanCan datasets. Based on these 4,873 KSAs, we created a ground-truth dataset that included 192,926 pairs of phosphosites known to be regulated by the same kinase (positives) and 1,956,116 pairs regulated by kinases from different kinase groups (negatives) (**Figure 1C, Supplementary Table 1**). These positive and negative site pairs include 2,410 unique sites, which we defined as annotated sites, and the sites not included in the ground-truth KSA dataset were referred to as unannotated sites. Since protein kinases are often either Serine-Threonine (S/T) kinases or Tyrosine (Y) kinases, we considered two types of pairs separately: S/T-S/T and Y-Y. Using this ground-truth KSA dataset, we classified the kinases into well-studied kinases (>10 substrates) and under-studied kinases (<=10 substrates).

As expected, compared to negative site pairs, positive site pairs had significantly higher 15 -mer peptide sequence similarities (**Figure 1D**), and their host proteins had significantly higher kinase interaction profile similarities in a protein-protein interaction network (**Figure 1E**). In addition, positive site pairs also showed significantly higher abundance correlations than negative pairs across all 11 cancer types (**Figure 1F**), largely independent of corresponding host protein correlations as demonstrated in heatmaps co-visualizing the distributions of log likelihood ratios (LLRs) between positive and negative pairs at the site (binned in rows) and protein (binned in columns) levels, respectively (**Figure 1G, Supplementary Figure 1A**). These data suggest that phosphosite co-expression, together with sequence similarity and host protein kinase interaction profile similarity, are effective predictors of co-regulation.

### CoPheeMap: a co-regulation map of the human cancer phosphoproteome

To create a network of co-regulated phosphosites, we developed an Extreme Gradient Boosting (XGBoost) classifier that integrates sequence similarity, host protein kinase interaction profile similarity, and pairwise phosphosite correlations for each of the 11 cancer types to distinguish the positive and negative phosphosite pairs (**Figure 2A**). This classifier was developed using carefully prepared training, validation, and independent test data based on the ground-truth dataset and 10 repetitions of Monte Carlo validation (**Supplementary Figure 2A-C, Methods**). The trained model incorporating all three types of features achieved an AUROC of 0.78 in the independent test data, outperforming models constructed using only subsets of features classified as static (sequence similarity scores and kinase interaction profile similarity scores) or dynamic (site-site correlations, **Figure 2B**). This performance gain remained evident when we applied a stringent False Positive Rate (FPR) threshold of 0.2% (**Figure 2B**).

**Figure 2:**
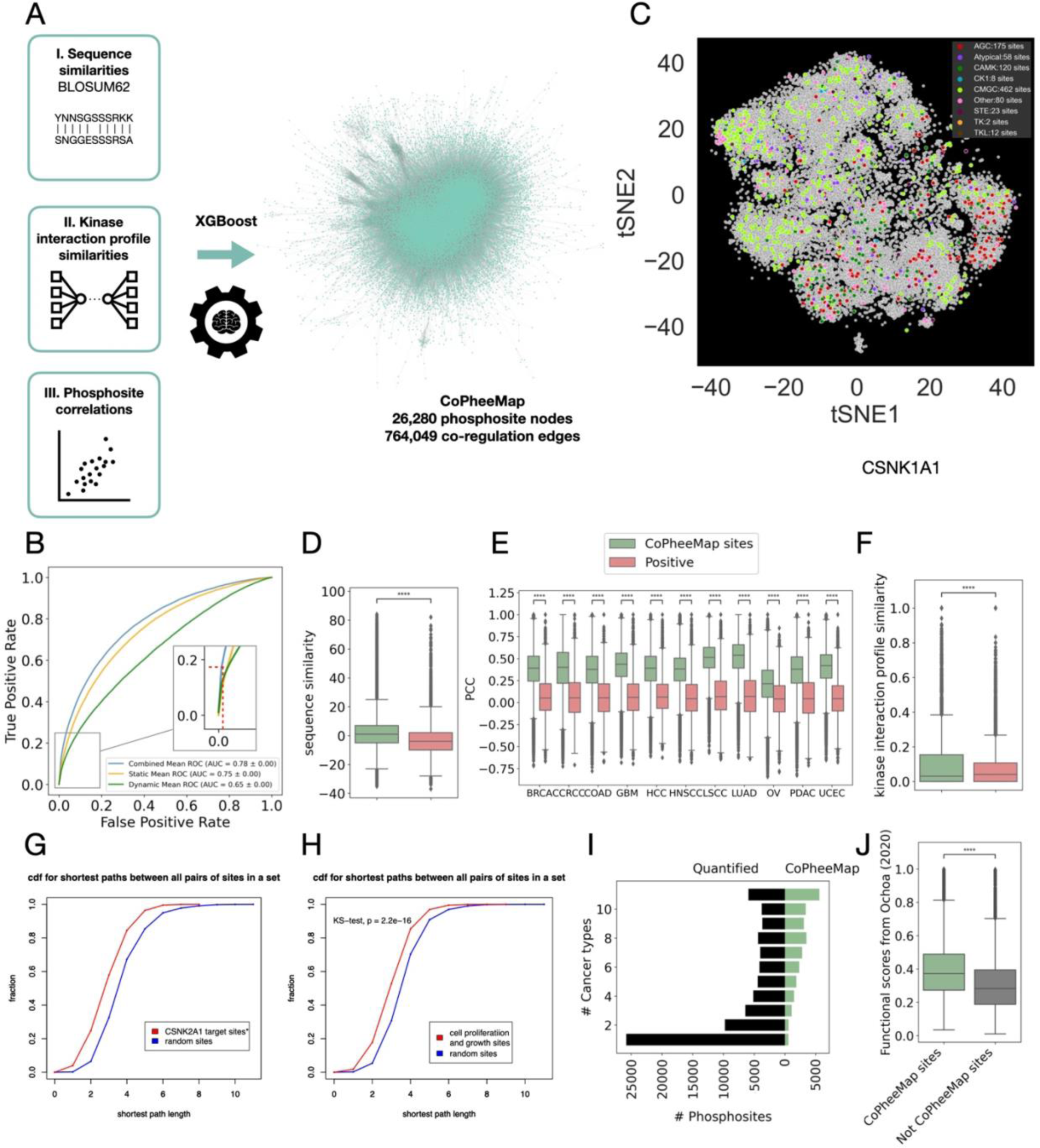
Construction and evaluation of the co-regulated network of phosphosites. A) Training, testing and application of the co-regulated phosphosite pair classifier (XGBoost model). The final CoPheeMap network contained more than 764K high-quality co-regulated phosphosite pairs constructed from 26,280 unique phosphosites. B) The model incorporating all three types of features achieved an AUROC of 0.78 in the independent test data, which was higher than models trained using only a subset of features. C) A 2-dimensional visualization of CoPheeMap utilizing Node2Vec encoding algorithms and tSNE. D-F) Comparison of sequence similarities (D), kinase interaction profile similarities (E) and abundance correlations (F) were significantly higher for the sites included in the CoPheeMap than for those from the ground truth positive set. G) Pairs of phosphosites regulated by one of the kinases excluded from the ground truth, CSNK2A1, were better connected than random pairs in the CoPheeMap. The empirical continuous distribution function (cdf) plots show that the distribution of shortest paths between all pairs of targets of CSNK2A1 was significantly lower than the distribution of a random set of pairs of sites of the same size. Ten CSKN2A1 targets in small networks unconnected to the main CoPheeMap network were excluded from this analysis. P-value from comparison of distributions using the Kolgmorov-Smirnov test was 2.23e-16. H) Pairs of phosphosites with related functional annotations were better connected in CoPheeMap than random pairs. The distribution of shortest paths between all pairs of sites associated by PhosphositePlus with the regulation of cell growth and proliferation was significantly lower than the distribution of a random set of pairs of sites of the same size. P-value from comparison of distributions using the Kolgmorov-Smirnov test was 2.23e-16. I) Phosphosites in CoPheeMap were more likely to be identified in multiple cancer types compared to those not included. J) Phosphosites in CoPheeMap also had higher functional significance scores from Ochoa et al. Nat. Biotchnol.

By applying the trained classifier to over 3 billion phosphosite pairs, controlling FPR at 0.2% to account for the large number of possible negatives, we identified 764,049 phosphosite pairs (0.03% of all candidate pairs) as co-regulated phosphosite pairs. These pairs, connecting 26,280 unique phosphosites, constituted a co-regulation network referred to as CoPheeMap (**Figure 2A, Supplementary Table 2**). To gain global insights into the organization of CoPheeMap, we generated 16-dimensional vector representations of all phosphosites in the network using the Node2Vec embedding algorithm. For visualization, we further reduced the representations to two dimensions using a multi-scale t-distributed stochastic neighbor embedding (tSNE) approach^30^ with parameters recommended for large network analysis^31^ (**Methods, Supplementary Table 2**). The distance between two phosphosites in the tSNE map reflects their distance in the coPheeMap network. In the tSNE map, phosphosites regulated by kinases in the same group tended to cluster together, further supporting the effectiveness of CoPheeMap in capturing co-regulation relationships between phosphosites (**Figure 2C**).

The newly predicted co-regulated phosphosite pairs in CoPheeMap even had significantly higher sequence similarity scores and co-expression levels than the positive pairs in the ground-truth data (**Figure 2D-E**). Although the median value of the kinase interaction profile similarity scores for the newly predicted co-regulated phosphosite pairs was relatively lower compared to the positive group, the newly predicted pairs were enriched with instances having high kinase interaction profile similarity scores, as demonstrated by the higher upper quartile (**Figure 2F**). Indicative of common targets of the same kinase being better connected than pairs of sites that are phosphorylated by different kinases, the positive site pairs had significantly shorter SPLs (shortest path lengths) than the negative site pairs did (**Supplementary Figure 2D**). Moreover, despite the sparsity of CoPheeMap, the known substrates of CSNK1A1, CSNK2A1, CSNK2A2, the three kinases intentionally excluded from the ‘ground truth’, were mostly well-connected (**Supplementary Figure 2E-G**). Particularly, the distribution of SPLs between all pairs of substrates of CSNK2A1, which had sufficient numbers for statistical assessment, was significantly lower than it was for pairs from a randomly drawn set of sites of the same size (KS test p=2.2e-16; **Figure 2G**), demonstrating the capacity of CoPheeMap to effectively recover missing co-regulation relationships within ground truth data.

Given that sites involved in the same biological function are more likely to be associated with common pathways regulated by common kinases, we reasoned that they would be closer together than random sites in the network. To test this, we collected functional annotations from PhosphoSitePlus for sites in CoPheeMap and parsed them into five categories of related functions: cell growth and proliferation, cellular degradation (apoptosis and autophagy), gene product regulation, microenvironment and mobility, and signaling pathway regulation. Consistent with our expectation, the distribution of SPLs between pairs of sites that are associated with the regulation of cell growth and proliferation is lower than that of pairs of sites from a randomly drawn set of the same size (KS test p=2.2e-16, **Figure 2H**). Similar results were obtained for all of the other functional categories (**Supplementary Figure 2H-K**), demonstrating that functionally related sites are closer together in CoPheeMap than randomly selected sites.

Further analysis showed that the phosphosites in CoPheeMap were more likely to be quantified in multiple cancer types compared with the other quantified phosphosites (**Figure 2I**). Sites included in CoPheeMap had higher abundance and higher predicted functional scores^32^ compared to those not included, suggesting high functional relevance for sites featured in CoPheeMap (**Supplementary Figure 2L, Figure 2J, Supplementary Table 2**). The degree distribution of CoPheeMap followed a power law (**Supplementary Figure 2M**). As expected, the annotated sites used for training tended to have higher degrees in the network compared to the unannotated sites; however, the latter also had a substantial number of connections in the network, with a median degree of 3.2, and some unannotated sites even acted as the hubs (**Supplementary Figure 2N**). For example, RFC1_S368, API5_S464, and TOP2A_S1106 had 1873, 1307, and 1283 connections, respectively (**Supplementary Table 2)**. In total, 98.6% of the edges (753,243 edges) in CoPheeMap involved at least one unannotated site. This extensive network of connections enabled the propagation of information from annotated to unannotated sites, an opportunity explored later in this paper.

### CoPheeKSA: a network-based generic KSA prediction model

Conventional methods for predicting KSAs typically rely on sequence motif information derived from known kinase targets^29, 33, 34^. These approaches are kinase-specific and lack generalizability when it comes to predicting KSAs for under-studied kinases (defined in this study as kinases with 10 or less known substrates). To develop a general KSA prediction model that could be applied to all kinases, we hypothesized that biological relationships among phosphosites, among kinases, and between kinases and phosphosites could all be utilized to improve generic KSA prediction.

To systematically capture biological relationships among phosphosites, we used CoPheeMap, which integrated both static and dynamic relationships. To capture biological relationships among kinases, we constructed a kinase network (KMap) where each edge represents co-expression at the protein level or physical interactions from a PPI network (**Methods, Figure 3A, Supplementary Table 3**). Applying the Node2Vec embedding algorithm to the two networks, we derived low-dimensional vector representations for 25,029 S/T phosphosites and 353 kinases (**Supplementary Table 2-3)**. These embeddings captured the inherent characteristics of each phosphosite and kinase in their network contexts.

**Figure 3:**
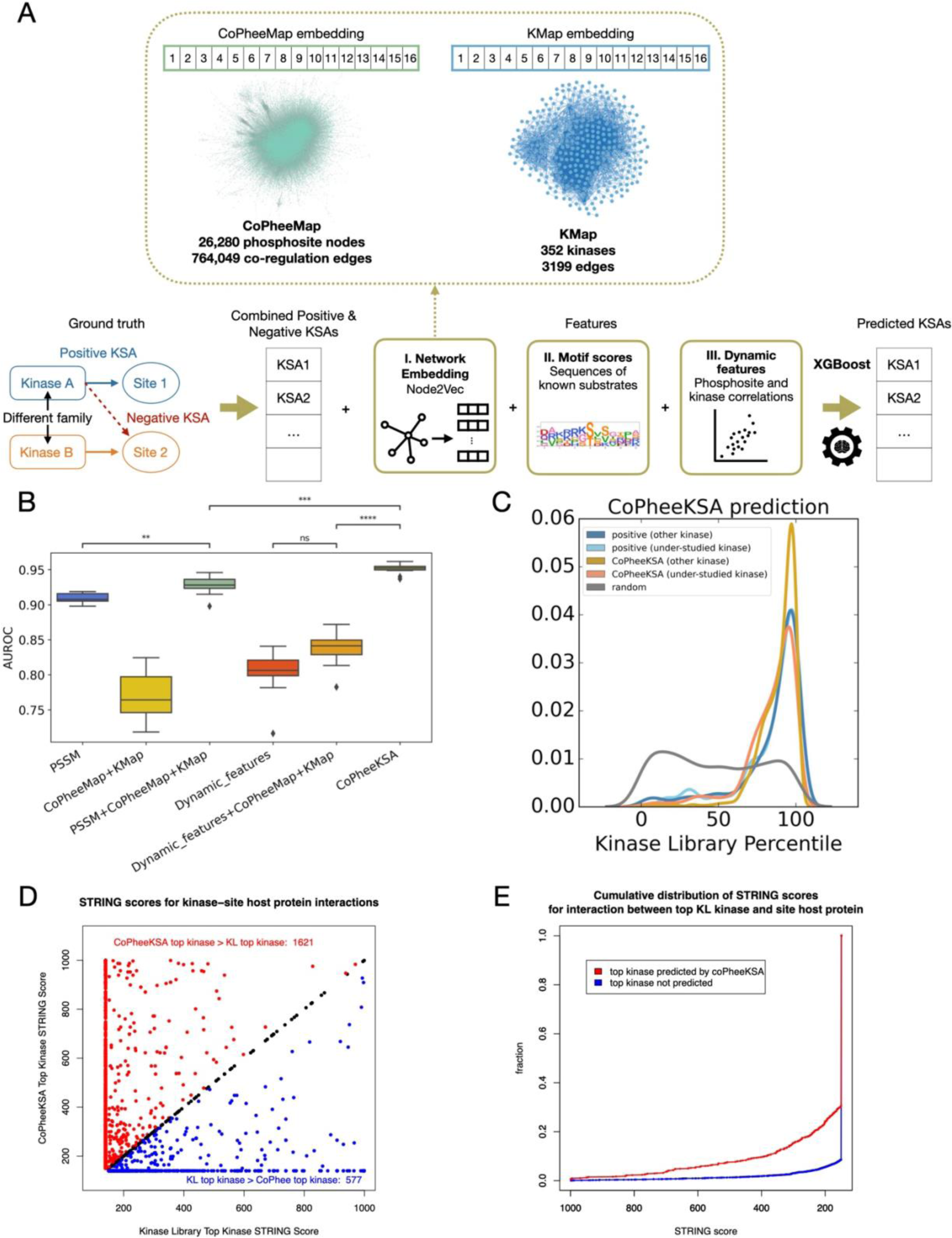
Development and evaluation of CoPheeKSA. A) Training, testing and application of network-based KSA classifier (XGBoost model), CoPheeKSA. To construct the ground truth positive KSAs for CoPheeKSA, we overlapped the substrates from the ground truth with CoPheeMap (Ser/Thr) using fifteenmer (+/-7 amino acids) and had 2,353 positive KSAs. To construct the negative KSAs for CoPheeKSA, we assigned known substrates of kinase A to kinase B if kinase A and B belong to different kinase families and both phosphorylate Ser/Thr. If phosphosites were annotated to be regulated by multiple kinases, the negative KSAs only included KSAs for which the site was a target of no overlapping up-stream kinases from the kinase groups for the kinases known to regulate the site. This approach yielded 114,530 ground truth negative KSAs. Heterogeneous features included network embeddings of CoPheeMap and Kmap, motif information, and dynamic features. KMap is a kinase network built upon PPI and the PanCan datasets. The links between kinases were defined as those with STRING PPI score > 400 or at least one protein-protein correlation > 0.5 in one cohort (proteomics data) among the 11 cancer types. B) AUROCs of KSA classifiers trained with different feature combinations. C) The kinase library percentile score distributions of the positive KSAs (under-studied kinases/well-studied kinases), the predicted positive KSAs (under-studied kinases/well-studied kinases) from CoPheeKSA and the random KSAs. The similar distributions between the positive and the predicted KSAs indicated good quality of those unique KSAs. D) Predictions from CoPheeKSA are more likely to reflect known protein-protein interactions between the kinase and substrate than are predictions from the kinase library (KL). For each site with any upstream kinases predicted by CoPheeKSA, the plot compares the STRING score for interaction of the host protein with the kinase with the highest prediction score from CoPheeKSA (y-axis) to that for the interaction with the kinase with the highest KL percentile score (x-axis). Black dots along the diagonal indicate sites where the top kinase predicted by CoPheeKSA was the same as the top kinase from the KL. Red dots above this line indicate sites where the STRING score was higher for the top kinase from CoPheeKSA than for the top kinase from the KL, while blue dots below the line indicate sites where the STRING score for the top KL kinase was higher than for the top CoPheeKSA kinase. E) The KSAs for the top kinases nominated by the KL for each site had higher STRING scores when the KSAs were also supported by CoPheeKSA than when they were not. The cdf plots show the distribution of STRING scores for interaction between the host protein of each site and the kinase with the highest percentile score for that site for KSAs predicted by CoPheeKSA (prediction score > 0.7676; red cdf) to that for KSAs not predicted by CoPheeKSA (prediction score < 0.7676; blue cdf). P-value from Kolgmorov-Smirnov test comparing the two distribution plots was 2.2e-16.

To capture relationships between each kinase and phosphosite pair, we first concatenated the corresponding kinase embedding from Kmap with the respective phosphosite embedding from CoPheeMap. Next, we computed position-specific scoring matrix (PSSM) scores for each kinase-phosphosite pair using the consensus motif of known substrates for that kinase (**Supplementary Table 3, Methods**). Moreover, since kinases typically interact with phosphorylation sites in a dynamic manner, utilizing proteomics and phosphoproteomics data, we further computed correlations between kinase protein abundance and phosphosite abundance, as well as between inferred kinase activity and phosphosite abundance, in each CPTAC cohort to capture their dynamic relationships (**Supplementary Table 3, Methods)**.

We used these features to train XGBoost models for KSA prediction (**Figure 3A**) based on a carefully prepared ground truth KSA dataset (**Supplementary Table 3, Methods**). Of note, to avoid information leakage, we computed kinase activity scores for each positive KSA separately, removing the corresponding phosphosite in our computation. The model integrating all features achieved a median AUROC of 0.95 in 10 repetitions of Monte Carlo validation, significantly outperforming PSSM-based prediction and XGBoost classifiers using only subsets of features (**Figure 3B, Supplementary Figure 3A**), suggesting that these features held complementary information. To ensure high quality predictions, we chose a highly stringent prediction threshold with an LLR of 5.5, i.e., the predicted KSAs were 244 times more likely to be positive KSAs than negative KSAs within the ground-truth data (**Supplementary Figure 3B**). With this stringent threshold, CoPheeKSA identified 24,015 KSAs involving 9,399 phosphosites and 104 S/T protein kinases, including 26 under-studied kinases (**Supplementary Table 3**).

To assess the quality of the CoPheeKSA predictions, we leveraged substrate specificities experimentally determined via the recently published Kinase Library (KL), where a higher KL percentile score corresponds to higher specificity^28^(**Supplementary Table 4).** We divided KSAs from ground truth and CoPheeKSA into two groups, those related to under-studied kinases and those linked to well-studied kinases. For the well-studied kinases, KSAs predicted by CoPheeKSA showed significantly higher KL percentile scores than the ground truth positive KSAs, (t-test, p<=1.00e-04, **Figure 3C, Supplementary Figure 3C**). For under-studied kinases, no significant difference was observed between the score distributions of the two groups (t-test, p > 0.01, **Figure 3C, Supplementary Figure 3C**). These results demonstrate the high quality of the CoPheeKSA predictions.

For comparison, we also investigated predicted upstream kinases for all the 77,442 quantified phosphosites in our PanCan data using a popular KSA prediction model, NetworKIN^34^. For both under-studied kinases and well-studied kinases, KSAs predicted by NetworKIN had significantly lower KL percentile scores than the ground truth KSAs (t-test, p<=1.00e-4, **Supplementary Figure 3D**). Furthermore, CoPheeKSA identified many more KSAs than NetworKIN (**Supplementary Figure 3E**). Notably, CoPheeKSA identifications covered more under-studied kinases (**Supplementary Figure 3F**). Given that these two KSA prediction approaches employ different methodologies utilized and have different phosphosite coverage scopes, it is not surprising that NetworKIN and CoPheeKSA made positive predictions for different phosphosite sets in the PanCan data (**Supplementary Figure 3G**).

We observed that many KSAs with high KL percentile scores received low scores in CoPheeKSA predictions (**Supplementary Figure 3H**). This discrepancy could be because KL scores represent a kinase’s potential to phosphorylate a substrate in vitro, which might not translate to actual interactions in vivo due to cellular conditions or protein structure constraints. To investigate if CoPheeKSA, trained with dynamic tumor data, more accurately predicts KSAs likely to occur in vivo, we used STRING scores for the interaction between the host protein of each site included in CoPheeKSA and the top kinase nominated to phosphorylate that site by CoPheeKSA (y-axis) or by the KL (x-axis) as a gauge for the likelihood of interaction between kinases and their putative substrates in a biological setting. Notably, for 1621 sites, the top kinase identified by CoPheeKSA had a higher STRING score than the one identified by the KL (red points above the black line in **Figure 3D**), while for only 577 sites, the KL’s top kinase scored higher (blue points). Further analysis showed that KSAs predicted by CoPheeKSA generally had higher STRING scores than those not supported by it (KS test p=2.2e-16; **Figure 3E**). Additionally, protein levels of kinases supported by CoPheeKSA correlated more strongly with their associated sites compared to unsupported kinases (KS test p=2.2e-16, **Supplementary Figure 3I**). Because the KL approach only captures peptide-based in vitro binding information, these data suggest that, by incorporating dynamic in vivo information in CoPheeMap and CoPheeKSA, our approach is able to discern between KSAs supported by cellular interactions from those that are not for high-scoring KSAs based on the KL approach and identify those that are more likely to manifest in vivo.

### Novel KSA predictions

To gain more insights into the newly predicted KSAs, we colored the phosphosites according to the up-stream kinases from prior knowledge and CoPheeKSA in the tSNE embeddings (**Figure 4A**). The co-clustering of the newly predicted substrates with previously annotated substrates supports the ability of CoPheeMap to connect phosphosites regulated by the same or similar kinases.

**Figure 4:**
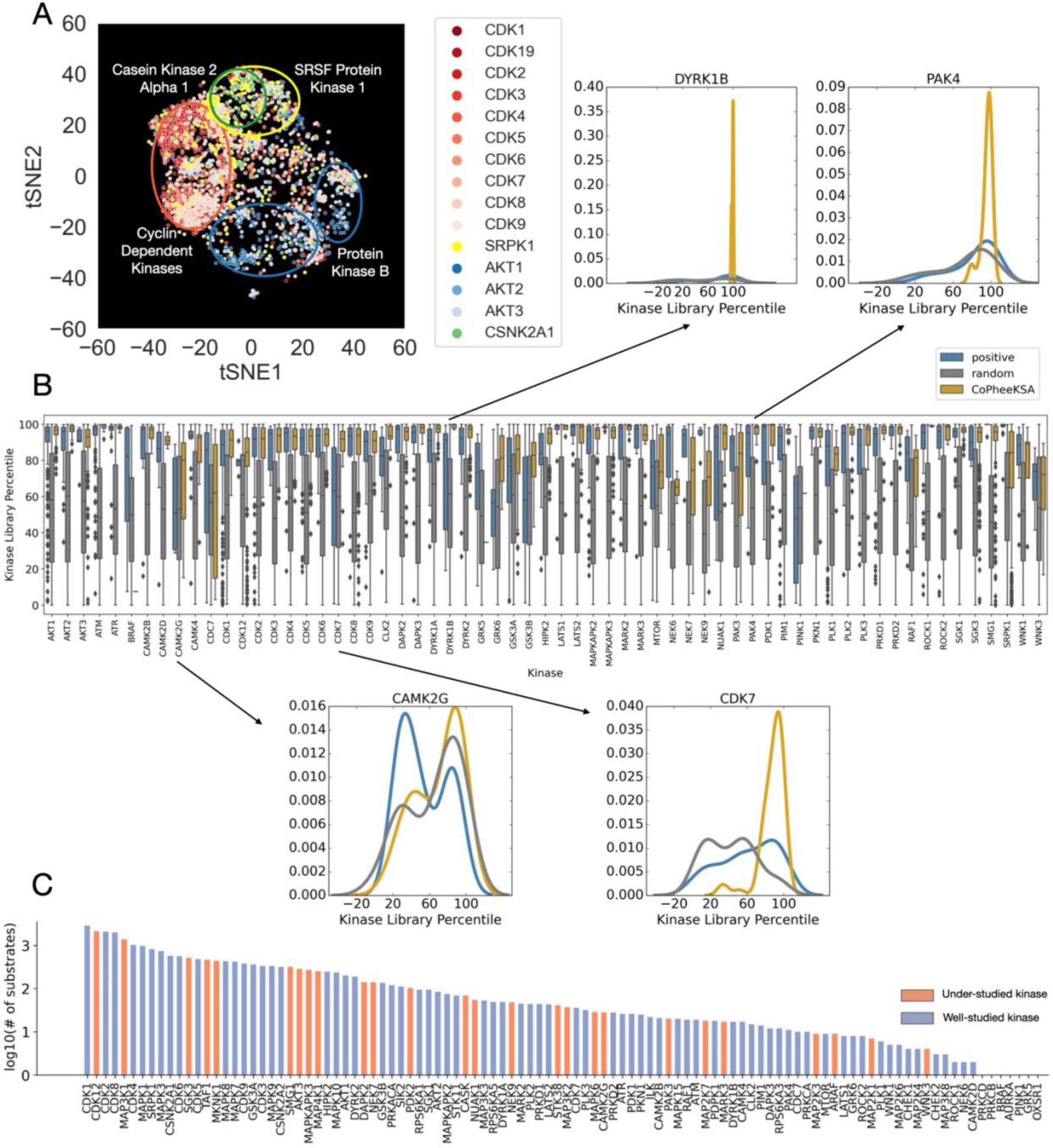
KSA predictions from CoPheeKSA. A) Two-dimensional visualization of CoPheeMap utilizing Node2Vec encoding algorithms and tSNE color coded by kinase for KSAs identified by CoPheeKSA combined with those included in the ground truth positive set. B) For individual kinases, the kinase library percentile score distributions of the positive KSAs (under-studied kinases/well-studied kinases), the predicted positive KSAs (under-studied kinases/well-studied kinases) from CoPheeKSA and the random KSAs. C) Number of substrates predicted for each kinase (under-studied and well-studied kinases are labeled).

For each kinase, we further compared the KL percentile score distributions between ground truth positive KSAs, random KSAs and predicted positives from CoPheeKSA. For most kinases, both ground truth positives and predicted positives had median KL percentiles over 90%, clearly higher than the KL percentile scores of the random group (**Figure 4B**). However, for some kinases, such as CAMK2G, CDK7, DYRK1B and GRK6, the median KL percentile scores for the ground truth positives were similar to random (**Figure 4B**). Interestingly, for these kinases, the predictions from CoPheeKSA had much better alignments with the KL data, evidenced by significantly improved KL percentile scores.

The number of novel KSAs predicted by CoPheeKSA for each kinase is visualized in **Figure 4C**. Not only did well-studied kinases like CDK8 and MAPK1 obtain more substrates, under-studied kinases, such as CDK12, SGK3, SMG1 and NUAK1, were also associated with hundreds of novel substrates. This could significantly enhance downstream analyses of under-studied kinases. Accordingly, we constructed a comprehensive KSA database integrating the KSA information from ground truth and CoPheeKSA (**Methods**). This KSA database includes 8,712 substrates, tripling the number of substrates in the ground truth data (2,844).

To confirm novel KSAs, we utilized IDPpub, a tool we recently developed for mining phosphosites from the abstracts of published papers^35^, and found 99 novel KSAs with potential support (**Supplementary Table 5**). Manual review of the corresponding evidence sentences confirmed 46 potential novel KSAs involving 20 unique kinases, including the 46 KSAs validated by evidence sentences from IDPpub confirmed novel substrates for 20 unique kinases, including 16 for CDK1, 5 for CDK2, and 4 for CDK5 as well as substrates for under-studied kinases. For example, the AKT3-dependent phosphorylation of FOXO3 at Ser253^36^ was not annotated in PhosphoSitePlus. Moreover, the MAP4K1/Hpk1-dependent phosphorylation of BLNK at Thr152^37^ was only noted in PhosphoSitePlus for the mouse ortholog, despite evidence showing that the phosphorylation of the endogenous human BLNK protein is also dependent on MAP4K1 in a human B cell line. This analysis shows CoPheeKSA can capture KSAs with support from the literature that were not previously identified by existing resources.

### Illuminating dark functional phosphosites

To illustrate the utility of CoPheeMap and CoPheeKSA for illuminating phosphosites of interest, we investigated phosphosites that had previously been prioritized by machine learning as functionally important sites, but without known up-stream kinases. The functional scores of the phosphosites were computed through machine learning-based integration of 59 features indicative of proteomic, structural, regulatory, or evolutionary relevance^32^ (**Supplementary Table 6**). Among the top 50 phosphosites in CoPheeMap with the highest functional scores, 39 had known up-stream kinases (**Figure 5**). The higher ratio of phosphosites with known up-stream kinases in this range was expected because of the intrinsic bias toward well-studied phosphosites in the ground truth dataset used to train the machine learning model for functional scoring^32^. Despite the bias, 11 out of the 50 did not have known up-stream kinases.

**Figure 5:**
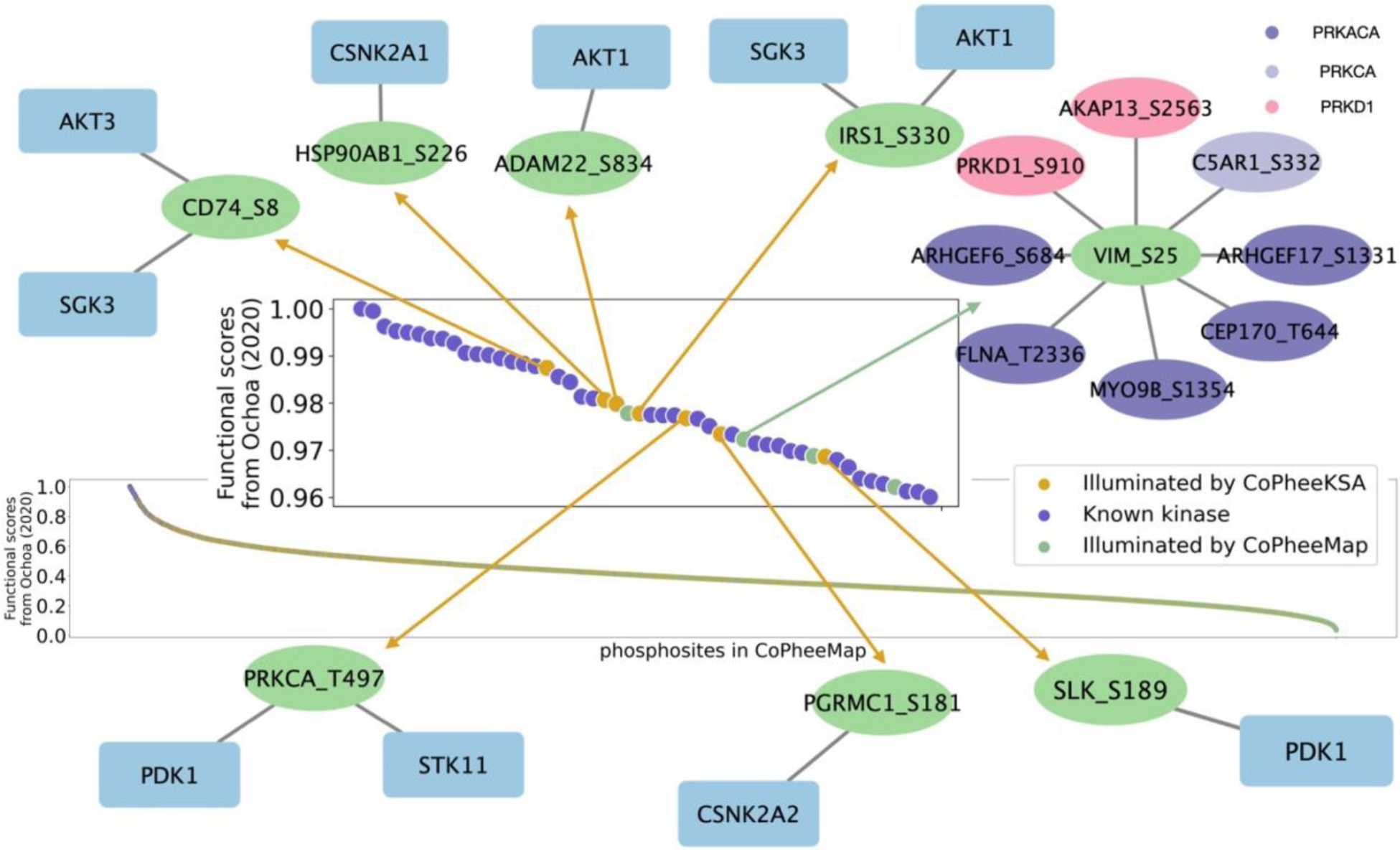
Illuminating phosphosites with top functional significance from Ochoa et al. Nat. Biotchnol. and the top differential phosphosites. Phosphosite with the top functional significance scores.

CoPheeKSA predicted up-stream kinases for seven of these 11 functional sites (**Figure 5**). One example is Ser8 on CD74, a cell-surface receptor and oncogene that has significantly higher protein abundance in tumors compared to normal tissues in multiple cancer types^7, 10, 11, 38^. RAC-gamma serine/threonine-protein kinase (AKT3) and serum/glucocorticoid regulated kinase family member 3 (SGK3) were predicted by CoPheeKSA as potential kinases regulating this functional site, and both kinases had KL percentile scores > 99% for this phosphosite^28^ (**Supplementary Table 5**). Consistent with the previously reported role for CD74 in immune stimulation^39^, AKT3 can be activated when immune cells are stimulated, performing essential functions in both innate and adaptive immune cells^40^.

As another example, Thr 497 is located in the activation loop of PRKCA, a Protein Kinase C family member that has been reported to play important roles in different cellular processes^41^ (**Figure 5**). For this phosphosite, Pyruvate Dehydrogenase Kinase 1 (PDK1) was the kinase predicted by CoPheeKSA, and PDK1 also ranked the first in the Kinase Library prediction (**Supplementary Table 6**). Although the database we used to generate ground truth data does not list any up-stream kinases for this site, PDK1 has been reported to phosphorylate Thr 497 on PRKCA, leading to secondary autophosphorylation events and conformational changes in the molecule^42^.

The third example is Ser 226 on heat shock protein 90 alpha family class B member 1 (HSP90AB1) (**Figure 5**). CoPheeKSA nominated Casein Kinase 2 Alpha 1 (CSNK2A1) as the regulatory kinase, which was also ranked top by Kinase Library (**Supplementary Table 6**).

Even for the remaining four dark phosphosites without confident CoPheeKSA predictions, CoPheeMap connected them to neighbors with known up-stream kinases or functional annotation, which could provide useful information for these sites because neighboring sites in CoPheeMap are likely to be co-regulated. For example, Ser 25 on VIM, a type III intermediate filament protein, is connected to substrates of PRKACA, PRKCA and PRKD1 in CoPheeMap. The regulatory kinases of these neighboring substrates were derived from both ground truth KSAs and CoPheeKSA predicted KSAs. Supporting this network neighborhood based inference, both PKC and PKA family members ranked at the top in the Kinase Library predictions for Ser 25 on VIM (**Supplementary Table 6**).

### Elucidating cancer-associated phosphosites and their regulatory kinases

To demonstrate the utility of CoPheeMap and CoPheeKSA in facilitating the analysis and interpretation of data from phosphoproteomics experiments, we conducted differential analysis of phosphosites between tumor and normal adjacent tissue (NAT) samples in each of the 8 CPTAC cohorts with both tumor and NAT samples and then calculated meta-p values to identify phosphosites that were differentially regulated across multiple cancer types.

First, we examined individual sites with the most significant differential abundance between tumor and NAT samples, focusing on those for which host protein differences were not nearly as significant (**Figure 6A, Supplementary Table 6, Methods**). Interestingly, several of these sites, including the most significant examples, Ser 67 on Nucleolin (NCL), Ser 153 on ESF1 nucleolar pre-rRNA processing protein homolog (ESF1), and Ser 254 on nucleophosmin 1 (NPM1), were predicted by CoPheeKSA to be regulated by cyclin dependent kinases, which have crucial roles in promoting the cell cycle. Abnormal activation of CDKs has been associated with tumor cell proliferation and cancer development^43^.

**Figure 6:**
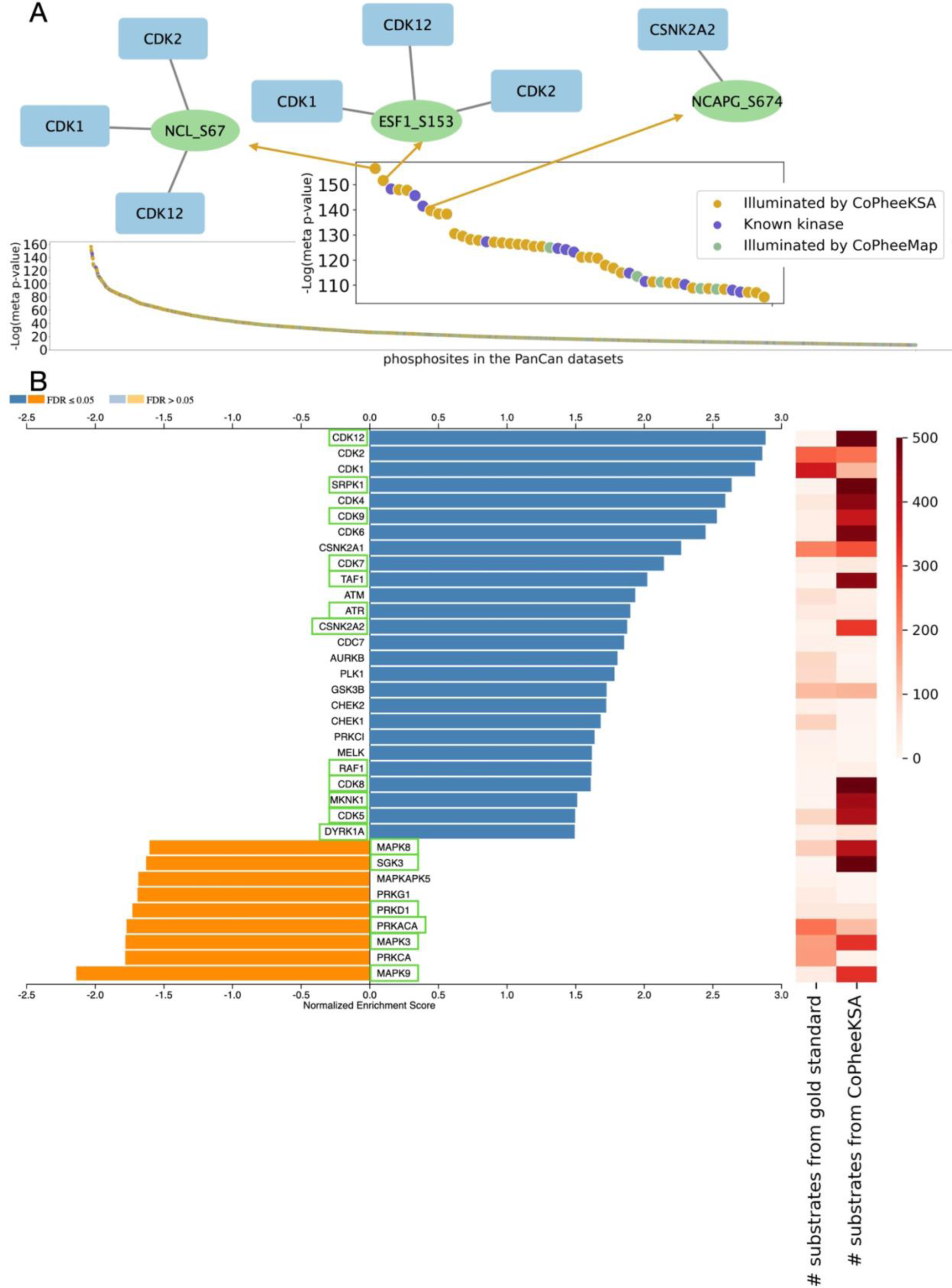
Hyper-active kinases identification in the PanCan datasets. A) Phosphosites with top -log10(meta p-value) in the differential analysis comparing tumor/normal. B) Up- and down-regulated kinases in the PanCan dataset from GSEA with the number of substrates from the ground truth and CoPheeKSA. The KSA database is a combination of ground truth and predictions from CoPheeKSA. The green labeled kinases are those uniquely identified using the comprehensive KSA database compared to the ground truth

To systematically identify hyperactivated kinases across cancer types, we used the meta-p values from our pan-cancer tumor versus normal comparisons as inputs for phosphosite set enrichment analysis based on our expanded KSA database. Consistent with our initial observations that many individual phosphosites highly up-regulated in the PanCan datasets were regulated by cyclin-dependent kinases, we confirmed the enrichment of CDK targets in this systematic analysis. While hyperactivation of CDK1, CDK2, CDK4, and CDK6 could be inferred without expanding the KSA database to include CoPheeKSA predictions, up-regulated activity for the less well-studied CDKs, including CDK12, CDK9, CDK7, CDK8, and CDK5, was only identifiable by adding targets predicted by CoPheeKSA. Using CDK12 as an example, among the 496 sites that contributed to the enrichment signal, 492 were uniquely associated with CDK12 by CoPheeKSA (**Figure 6B, Supplementary Figure 5A**). Although the two previously curated target sites of this kinase were up-regulated in cancer samples, without CoPheeKSA predictions, they were insufficient to produce statistically significant enrichment in phosphosite set enrichment analysis. Consistent with the substrate-based hyperactivation inference for CDK12 and other less well-studied CDKs, they all showed elevated protein abundance across cancer types (**Figure 6B, Supplementary Figure 5B-F**).

In addition to these less well-studied CDKs, our analysis also identified well-studied kinases that would have not been identified without CoPheeKSA predictions (**FDR <0.05, Figure 6B, Supplementary Table 6**). These included serine-arginine protein kinase 1 (SRPK1), TATA-box binding protein associated factor 1 (TAF1), Raf-1 proto-oncogene, serine/threonine kinase (RAF1), MAPK interacting serine/threonine kinase 1 (MKNK1), dual specificity tyrosine phosphorylation regulated kinase 1A (DYRK1A), and several mitogen-activated protein kinases (MAPKs). Many of these inferences were also supported by elevated mRNA and protein abundance of the kinases in tumors compared to normal samples (**Supplementary Figure 5G-L**). Moreover, some of these kinases, such as SRPK1, have been previously associated with cancer staging, subtyping and survival^44^.

## Discussion

The ‘guilt-by-association’ strategy is commonly used to predict functions of under-studied genes and proteins^25, 45, 46^. Leveraging the vast amount of CPTAC pan-cancer phosphoproteomics data, our study expands this strategy to address the dark phosphoproteome challenge through the construction of CoPheeMap, a machine-learned co-regulation network of the human phosphoproteome. In conjunction with CoPheeKSA, which predicts KSAs by utilizing information from CoPheeMap, these new tools collectively offer a comprehensive framework for investigating the regulation and functions of phosphosites, particularly in the context of human cancer.

We showed that phosphosite co-expression is an effective predictor of their co-regulation relationship. Importantly, phosphosite co-expression, as well as neighboring sequence similarity and host protein kinase interaction profile similarity, the other two features we used to predict phosphosite co-regulation, are purely data driven and not affected by existing knowledge on their regulatory kinases. Built upon these features, CoPheeMap was able to include 26,280 phosphosites without prior functional or regulatory information. This also allowed CoPheeKSA to make predictions that covered more under-studied kinases than other computational tools such as NetworKIN. The high quality of the CoPheeKSA predictions was validated using independent experimental data from the Kinase Library as well as the literature mining tool IDPpub. Interestingly, for some kinases, new predictions from CoPheKSA aligned much better with the KL scores than the ground truth positives, suggesting that our generic model may overcome noise in the ground truth data. We also found that many KSAs with high scores in the Kinase Library had low scores in CoPheeKSA, suggesting that CoPheeKSA could be used to refine predictions from KL.

Although we can now generate comprehensive measurements of protein phosphorylation events, our understanding of phosphoproteomics data is limited by insufficient knowledge about the regulatory and functional aspects of phosphosites. By utilizing computationally predicted functional sites from a published study^32^ and cancer-associated phosphosites identified through a pan-cancer analysis, we demonstrated how CoPheeMap and CoPheeKSA can help uncover regulatory and functional insights into biologically significant yet underexplored phosphosites. Our analysis of the cancer-associated phosphosites revealed multiple hyperactivated kinases that have been largely overlooked but could serve as potential therapeutic targets. For example, CDK12, a kinase implicated in the regulation of MYC expression, Wnt/β-catenin signaling, RNA splicing, and ErbB-PI3K-AKT signaling^47^, was identified as a candidate. CDK12 inhibitors have been shown to suppress cancer cell transcription and growth and enhance drug susceptibility^47–50^. The inhibition of CDK12 has demonstrated promising therapeutic effects on treating cancers, particularly those driven by dysregulated transcription factors^47, 48^. Our findings suggest that further research into CDK12 inhibitors in cancer treatment is warranted and highlight potential phosphosite biomarkers for monitoring CKD12 activity and the effectiveness of its inhibitors.

We identified several areas for improvement in future research. Firstly, during the construction of the ground truth of co-regulated phosphosite pairs for the construction of CoPheeMap, we found that certain kinases can phosphorylate both serine and tyrosine residues. This occurrence, though rare, introduces noise that may affect the training of machine learning models. Therefore, we limited our dataset to pairs of either Ser/Thr-Ser/Thr or Tyr-Tyr sites. Due to the scarcity of tyrosine data from the experimental methods used by CPTAC, particularly across multiple cancers, the phospho-tyrosine subnetwork in CoPheeMap includes only about 300 tyrosines. Consequently, CoPheeKSA was not applied to this subnetwork, limiting insights into tyrosine kinases, which are crucial targets for oncology and therapy development. We aim to enrich future studies with tyrosine-enriched data to enhance CoPheeMap’s coverage. Additionally, in our construction of the CoPheeKSA models, the requirement for a kinase to have at least 5 known substrates to be included left some serine/threonine kinases unexplored. Integrating Kinase Library data into our model training could be a significant enhancement.

Secondly, to minimize bias towards well-researched kinases, CoPheeMap focuses only on co-regulated site pairs without considering specific kinase information. This strategy allowed CoPheeMap to include a substantial number of unannotated phosphosites, and some even acted as the hubs. However, sites regulated by well-known kinases are more likely to be the hubs in the network. This intrinsic bias in the training data is difficult to avoid because it is already present in the KSA databases used for ground truth construction. The use of the unbiased Kinase Library data may provide a solution to address this bias.

Finally, in our study, we utilized data from 11 CPTAC cancer types to construct the pan-cancer CoPheeMap. As additional phosphoproteomics data for other cancer types become available, we plan to incorporate them in our network construction process to improve the network’s comprehensiveness. Our approach can also be used to integrate phosphoproteomics datasets from a single cancer type, allowing for the creation of cancer type-specific CoPheeMaps. Furthermore, this strategy extends beyond cancer to enable the construction of disease-specific networks for various diseases, offering wide-ranging applications in biomedical research.

## Methods

### Data acquisition

CPTAC clinical data were downloaded from the CPTAC Data Portal and https://pdc.cancer.gov/pdc/cptac-pancancer as described in Liao et al. The clinical data used in the portal were collected from CPTAC with the May 2022 update. Age was truncated to 90 years. Tumors with a size <=0 were replaced with NA.

CPTAC RNA-seq sequence data and proteomics data were downloaded from NCI Genomics Data Commons (GDC) and Proteomics Data Commons (PDC) as described in Liao et al. All sample IDs were mapped to the patient Case ID and replicate samples were discarded. Cases and samples not used in the flagship manuscripts^10–14, 38, 51–54^ were also discarded.

### RNA data processing and quantification

We obtained the gene-level read count and Fragments Per Kilobase of transcript per Million mapped reads (FPKM) using RSEM v1.3.1^55^ and Bowtie2 v2.3.3^56^. We used a common reference genome annotation, GENCODE V34 basic (CHR)^57^. We normalized the upper quantiles of coding gene RSEMs to 1500. Then, the normalized RSEM values were log2-transformed.

### Proteomics and phosphoproteomics data processing

MSFragger v3.4^58^, the Philosopher v4.0.1^59^ toolkit, and the TMT-Integrator^60^ pipeline were used by the Michigan University team in the CPTAC pan-cancer working group to process and quantify the mass spectrometry data as described in Liao et al.. A single primary isoform was selected for each gene. For coding genes, MANE Select and SwissProt were used to prioritize isoforms. If a gene did not have a single MANE Select and/or SwissProt isoform, then the isoform was prioritized using the longest protein sequence followed by the longest transcript. Additionally, remaining Swiss-Prot proteins and MANE Plus Clinical isoforms were retained as secondary isoforms for web portal display.

We normalized data with median centering the medians of the reference intensities. Proteins/phosphosites that were quantified in at least 20% of samples in at least one cohort were included in downstream analysis.

### Phosphoproteomics isoform mapping

We used both primary and secondary protein isoforms to re-annotated phosphosites as described in Liao et al. The site ID finally consisted of the Ensembl gene ID, Ensembl protein ID, site position based on the selected protein ID, fifteenmer (+/- 7 amino acids) based on the selected protein ID, and a flag for whether the protein is a primary (1) or secondary (2) selected sequence.

### Ground truth dataset of CoPheeMap

The ground truth kinase substrate associations were downloaded from GPS 5.0, http://gps.biocuckoo.cn, which is a curation of 15,194 experimentally identified kinase substrate associations (**Supplementary Table 1**). To classify the kinases, we further annotated the kinases with kinase group information from KinBase, http://kinase.com/web/current/kinbase.

The phosphosite pairs regulated by the same protein kinases were defined as the positive pairs. The phosphosite pairs regulated by the kinases from different kinase groups were defined as the negative pairs. We used fifteenmer (+/- 7 amino acids) to represent the positive and negative phosphosite pairs (**Supplementary Table 1**). If phosphosites were annotated to be regulated by multiple kinases, no overlapping known up-stream kinase families were allowed in the negative pairs. Only Ser/Thr-Ser/Thr (S/T-S/T) pairs or Tyr-Tyr (Y-Y) pairs were kept.

### Sequence similarity scores

To calculate phosphopeptide sequence similarities, we used the BLOSUM62^61^ matrix. The similarity score between the pair of fifteenmers (+/- 7 amino acids) of phosphosite A and phosphosite B was defined as the sum of the log-odds ratios from BLOSUM62 for all flanking positions (−7 to −1 and +1 to +7) (**Supplementary Table 1**).

### Kinase interaction profile similarity scores

We downloaded the STRING PPI network (protein.links.v11.5) and kept the links with scores > 400. For each phosphosite, we defined the list of kinases linked to the host protein as those with PPI scores > 400 for the pair. We then calculated the Jaccard index of the two kinase lists as the kinase interaction profile similarity scores between each pair of phosphosites (**Supplementary Table 1**).

### Phosphosite/host protein abundance correlations and likelihood ratios

For each phosphosite pair or protein pair, we require quantification in at least 20 overlapping samples in one cohort for calculating the Spearman’s correlation coefficient (**Supplementary Table 1**).

We then divided all the phosphosite pairs into 361 groups which we also called bins (19*19). In each bin, the pairs had site-site/protein-protein abundance correlations ranging from −0.9 to 1 with an interception of 0.1 and matched host protein-host protein abundance correlations ranging from −0.9 to 1 with an interception of 0.1. The log likelihood ratios (LLRs) of co-regulation were defined as the fraction of ‘ground-truth’ positive site pairs divided by the fraction of negative site pairs in the each one of the total 361 bins (19*19 bins).

### Training, evaluation, test data and prediction data for CoPheeMap

To avoid overfitting, we separated the ground-truth phosphosites into training, evaluation and test groups (**Supplementary Figure 2A**), ensuring that no phosphosites overlapped between these groups. Links between the training sites and the evaluation sites were assigned to the evaluation data. Links between the training sites and the test sites or the links between the evaluation sites and the test sites were assigned to the test data. The ratio between training + evaluation and test sites or between training and evaluation sites for Ser/Thr is 9:1. This ratio for Tyr is 5:5 due to the smaller sample size. The ratios of positive and negative pairs in the training, validation and test data were 1:5, 1:5 and 1:50 (**Supplementary Figure 2B**). More than 3,000,000,000 phosphosite pairs had at least one site-level correlation calculated in one or more cohorts. All site pairs were prepared as the prediction data. The three kinases excluded from ground truth were CSNK1A1, CSNK2A1 and CSNK2A2.

### Extreme Gradient Boosting (XGBoost) classifier for CoPheeMap

XGBoost was trained on the ground truth data to generate models to classify each pair of sites as being co-regulated by the same upstream kinase or not. For each pair of sites, the 14 input features including 11 for pairwise phosphosite correlations from each of the 11 cohorts, 1 for sequence similarity scores, 1 for kinase interaction profile similarity scores and 1 label indicating Ser/Thr-Ser/Thr or Tyr-Tyr (**Supplementary Table 1**). Missing values were kept. We fine-tuned the XGBoost hyper parameters using the evaluation data. Different classifiers were trained using all the 14 features, the 11 dynamic features or the 2 static features. AUROCs were calculated for each classifier using test data.

### Network Embedding and dimensionality reduction

The networks (CoPheeMap and KMap) were embedded with Node2Vec^62^ with dimensions=16, window=1, min_count=1, batch_words=4 (**Supplementary Table 2**). Default tSNE was used to reduce the dimensionality of CoPheeMap embedding features from 16 to 2 (**Supplementary Table 2**).

### Network visualization

The networks were directly visualized using Cytoscape 3.9.1 or Gephi 0.10.1.

### Shortest Path Lengths (SPLs)

The iGraph R (networkx Python) package was used to calculate the shortest path length between each pair of nodes in CoPheeMap that belong to the indicated group (i.e., targets of CSKN2A1 or sites annotated as regulating the same functional category) or a set of random pairs of the same size.

### Analysis of functionally annotated sites

We downloaded the Regulatory_sites database file from PhosphositePlus (phosphosite.org) on October 26, 2023. The “ON_PROCESS” column in this file includes information about cellular processes that are annotated as being regulated by specific phosphosites. After filtering the sites in this file to those that matched the host protein HGNC symbols and fifteenmers of the sites included in the CoPheeMap embedding, sites were grouped into 5 selected categories with sufficient numbers of sites associated with functionally distinct cellular processes included in the network (>100). These groupings included cell growth and proliferation (all ON_PROCESS categories including “cell growth” or “cell cycle regulation”, regardless of direction of regulation), gene product regulation (ON_PROCESS categories including the key words “transcription”, “translation”, “chromatin organization”, “RNA splicing”, or “RNA stability”), cellular degradation (ON_PROCESS categories including “apoptosis” or “autophagy”), tumor microenvironment and mobility (ON_PROCESS categories including “cytoskeletal reorganization” or “cell motility”), and signal transduction (ON_PROCESS categories including “signaling”).

### Phosphosite functional scores

The functional scores were from the functional landscape of human phosphoproteome^32^. The sites were mapped using uniprot ids and site positions (**Supplementary Table 2**).

### Kinase network (KMap)

The links between kinases were defined as the STRING PPI score > 400 or at least one protein-protein correlation > 0.5 in one cohort (CPTAC proteomics data) among the 11 cancer types. This network covered 352 kinases with 3,199 edges (**Supplementary Table 3**).

### Ground truth dataset of CoPheeKSA

Based on the ground truth kinase substrate associations, we classified kinases as under-studied kinases (<= 10 known substrates) and well-studied kinases (> 10 known substrates).

To construct the positive KSAs for CoPheeKSA, we overlapped the substrates from the ground truth with CoPheeMap (Ser/Thr) using fifteenmer (+/- 7 amino acids) and had 2,353 positive KSAs. To construct the negative KSAs for CoPheeKSA, we assigned known substrates of kinase A to kinase B if kinase A and B belong to different kinase families and both phosphorylate Ser/Thr. If phosphosites were annotated to be regulated by multiple kinases, the negative KSAs only included KSAs for which the site was a target of no overlapping up-stream kinases from the kinase groups for the kinases known to regulate the site. This approach yielded 114,530 negative KSAs (**Supplementary Table 3**).

### Kinase activity scores

The kinase activity scores were the mean values of all known substrates from the ground truth quantified in that cancer cohort. For each kinase activity score, we required quantification of at least 3 substrates. To avoid overfitting, for each KSA in the ground truth where the target site is on the kinase itself?, the site abundance was removed when calculating the kinase activity score for the corresponding kinase (**Supplementary Table 3**).

### Kinase motif scores

There is one static feature of motif scores for each KSA. The motif information (the sum of the possibilities for each amino acid in each position) was gained from the ground truth (a kinase should have at least 5 substrates to calculate the motif). To avoid overfitting, the test sites were removed when calculating the motif information for the test sites (**Supplementary Table 3**).

### Correlations between kinase and substrate

For correlations between kinase protein abundance/kinase activity scores and phosphorylation abundance, we require quantification in at least 20 overlapping samples in one cohort for calculating the Spearmans’ correlation coefficient (**Supplementary Table 3**).

### Training, evaluation, test data and prediction data for CoPheeKSA

The KSAs from different kinases were combined together as positive KSAs and negative KSAs. We separated the positive KSAs into training and test groups for Monte Carlo cross validation (10 times). 235 positive KSAs were included in each test group. The test groups were then split into evaluation and test groups with 25 positive KSAs in each evaluation group. To avoid the impact of missing values, we kept the same ratio of missing values for the dynamic features. All the potential kinase-substrate interactions were prepared as the prediction data.

### Extreme Gradient Boosting (XGBoost) classifier for CoPheeKSA

XGBoost was trained using the ground truth KSA data to generate models to classify each potential KSA (kinase-site pair) as a positive or negative KSA. For each KSA, the input included 55 features (16 features for CoPheeMap network embedding information from Node2vec, 16 features for KMap network embedding information, 22 dynamic features of correlations between the kinase and substrate in the KSA, and 1 static feature for similarity of the site sequence to the PSSM score for the kinase motif, **Supplementary Table 3**). We kept missing values for dynamic features. Because for each KSA in the ground truth, we removed the site abundance when calculating the kinase activity scores and led to a higher ratio of missing values in the kinase activity scores for the KSAs in the ground truth positive set, there is a higher ratio of missing values in the correlations between kinase activities and phosphorylation abundance in the ground truth positive set data, compared to the negative set data or the data for the predicted KSAs. To avoid overfitting, we intentionally and randomly removed some dynamic features (correlations between kinase activity scores and phosphorylation abundance) in the negative data to keep the same ratio of missing values as the positive KSAs. The ratio of the positives and the negatives in the training, evaluation and test data is 1:10.

Different classifiers were trained using all the features, only the motif scores, only the network embedding features, the network embedding features and the motif scores, only the dynamic features or the network embedding features and the dynamic features. AUROCs were calculated for each classifier (**Figure 3B**). The XGBoost parameter is {’max_depth’: 2, ‘eta’: 0.2, ‘objective’: ‘binary:logistic’, ‘num_round’: 300} with AUC as the evaluation metric. CoPheeKSA had a general threshold of 0.7676 with LLR=5.5 (**Supplementary Figure 3B**).

### The Kinase Library scores

Percentile scores for each phosphosite and kinase were computed as described in Johnson, et al^28^. Briefly, all phosphosites in this study were scored by all the characterized kinases (303 S/T kinases), and their ranks in an a-priori score distribution based on curated phosphoproteome were determined to yield percentile-score.

### NetworKIN prediction

We used the webtool of NetworKIN^34^ to predict the up-stream kinases for all the phosphosites quantified in the PanCan datasets. The threshold is 5.

### Comparison of STRING PPI Scores for CoPheeKSA and KL KSAs

To test whether training CoPheeKSA with the dynamic features from tumor data better captures KSA relationships that are more likely to occur in vivo than predictions from the kinase library does, we considered STRING scores for the interaction between site host proteins and kinases as a proxy for the likelihood that KSAs are biologically relevant. For this analysis, we used scores from the 9606.protein.links.v12.0.txt file from STRING (string-db.org) to assess PPI strength for the interaction between the host protein of each site with a predicted KSA by CoPheeKSA (prediction score > 0.7676) and the top scoring kinase from CoPheeKSA (kinase with highest prediction score for that site; when the same site is predicted to be the target of multiple kinases, only the best scoring kinase was used for the analysis in **Figure 3E**) or the top scoring kinase from the KL (kinase with highest percentile score for that site). Since STRING only provided scores for interactions with a minimum value of 150, any interaction not included in the STRING dataset was assumed to have little or no support and set to 149 unless scores were not available for the best kinase nominated by both CoPheeKSA and the KL, in which case the site was excluded. For the analysis in Figure 3F, all KSAs with no STRING score were set to 149.

### IDPpub validation

Evidence sentence data from IDPpub^35^ is available for download at https://www.zhang-lab.org/idppub/. We first searched the main results from IDPpub for exact substrate HGNC symbol and site amino acid position matches for each site with KSAs predicted by CoPheeKSA. We then extracted the evidence sentences for each matched site and searched the kinases_for_idppub.txt file for matches with the corresponding kinase from the predicted KSA. This file lists any kinases that were mentioned in the corresponding evidence sentence for each of the identified sites. However, IDPpub was trained only to identify phosphosites not KSAs, so the number of KSAs that can be confirmed using this approach is limited and required further manual review to validate the relationship between the kinase and the site. In order to focus on potential novel KSAs prior to manual review, any KSAs that were included in the ground truth positive set for CoPheeKSA were filtered out. If we assume that any KSA with support from more than 10 evidence sentences is a misannotation of a known KSA, this search yielded 99 novel KSAs with potential support from IDPpub (**Supplementary Table 5** also includes those with support from more than 10 sentences). Manual review of the corresponding evidence sentences directly validated 46 of the 99 potential novel KSAs with support from IDPpub where one of the corresponding evidence sentences specifically stated that the site from the KSA was phosphorylated by the kinase from the KSA.

### KSA database

We constructed the gmt files for GSEA/ssGSEA analysis. When a kinase has more than 100 known substrates, no predicted substrates were added. When kinase has less than 100 known substrates, the predicted substrates from CoPheeKSA were added up to the maximum number of 500 (ranked by predicted scores).

### Tumor versus normal comparison

Paired tumor samples and normal samples derived from 8 cancer types (CCRCC, COAD, HNSCC, LSCC, LUAD, OV, PDAC, and UCEC) for both proteomics and phosphoproteomics were used for differential expression analysis. Proteins were required to be detected in at least 20 tumor samples and 10 normal samples for proteomics and phosphoproteomics datasets. The unpaired Wilcoxon Rank Sum test was used to calculate significance.

### Gene set enrichment analysis (GSEA)

We used Webgestalt (http://www.webgestalt.org/) to do the GSEA on the phosphosites and the - log10(p-values)^63^. The organism was set to ‘other’. The functional database was uploaded using self-defined gmt files.

### Hallmark pathway single sample gene set enrichment analysis (ssGSEA)

ssGSEA was performed for each cancer type using gene-wise Z-scores of the RNA expression data (RSEM) for the MSigDB Hallmark gene sets v7.0^64^ via the ssGSEA2.0 R package^65^. RNA data were filtered to coding genes with < 50% 0 expression. (Parameters: sample.norm.type=”rank”, weight=0.75, statistic=”area.under.RES”, nperm=1000, min.overlap=10). Pathway activity scores are normalized enrichment scores from ssGSEA.

### Statistical associations

For all association tests, at least 10 samples within a group were required to have measurements. The statistical test was tailored to the data type. Spearman’s correlation was used for continuous data, Jonckheere-Terpstra trend test for ordinal data, Student’s T-test for binary data, and Cox regression for time to event data. For the enrichment analysis, we used Fisher’s Exact Test. In all the box plots, Student’s T-test independent samples with Bonferroni correction were applied for the distribution comparison. The Kolgmorov-Smirnov test was used to assess differences between distributions shown in empirical continuous distribution frequency (cdf) plots.

### Meta p-value calculation

Meta p-values were calculated with the “sumz” method from the R package metap (V1.4). P-values of individual cohorts were first converted to one-sided p-values and the sign for p-values not consistent with the majority were reversed. The calculated meta p-value was converted back to two-sided p-values and then the major sign of association was added.

## Acknowledgments

We gratefully acknowledge contributions from the CPTAC and its Pan-Cancer Analysis Working Group. This work was supported by National Institutes of Health (NIH) grants from the National Cancer Institute (NCI) U24 CA271076, R01 CA245903, and U01 CA271247, by Cancer Prevention & Research Institute of Texas (CPRIT) Award RP220050 and RP210027 - Baylor College of Medicine Comprehensive Cancer Training Program, and funding from the McNair Medical Institute at The Robert and Janice McNair Foundation. B.Z. is a CPRIT Scholar in Cancer Research and a McNair Scholar.

## Author Contributions

Conceptualization, B.Z., W.J.; Methodology, W.J.; Formal Analysis, W.J., E.J.J., T.M.Y., J.L.J.; Investigation, W.J., E.J.J., B.Z.; Resources, Y.L., T.M.Y., J.L.J., L.C.C.; Data Curation, W.J.; Writing - Original Draft, W.J., E.J.J., B.Z.; Visualization, W.J., E.J.J.; Supervision, B.Z., L.C.C.; Funding Acquisition, B.Z.

